# The SARS-CoV-2 receptor-binding domain elicits a potent neutralizing response without antibody-dependent enhancement

**DOI:** 10.1101/2020.04.10.036418

**Authors:** Brian D. Quinlan, Huihui Mou, Lizhou Zhang, Yan Guo, Wenhui He, Amrita Ojha, Mark S. Parcells, Guangxiang Luo, Wenhui Li, Guocai Zhong, Hyeryun Choe, Michael Farzan

**Affiliations:** Department of Immunology and Microbiology, The Scripps Research Institute, Jupiter, FL 33458, USA; Department of Animal and Food Sciences, University of Delaware, Newark, DE 19716, USA; Department of Microbiology, University of Alabama at Birmingham School Of Medicine, Birmingham, AL 35294, USA; National Institute of Biological Sciences, Tsinghua Institute of Multidisciplinary Biomedical Research, Tsinghua University, Beijing, China; Scripps Research | SZBL Chemical Biology Institute, Shenzhen Bay Laboratory (SZBL), Shenzhen, China; School of Chemical Biology and Biotechnology, Peking University Shenzhen Graduate School, Shenzhen, China

## Abstract

The SARS-coronavirus 2 (SARS-CoV-2) spike (S) protein mediates entry of SARS-CoV-2 into cells expressing the angiotensin-converting enzyme 2 (ACE2). The S protein engages ACE2 through its receptor-binding domain (RBD), an independently folded 197-amino acid fragment of the 1273-amino acid S-protein protomer. Antibodies to the RBD domain of SARS-CoV (SARS-CoV-1), a closely related coronavirus which emerged in 2002-2003, have been shown to potently neutralize SARS-CoV-1 S-protein-mediated entry, and the presence of anti-RBD antibodies correlates with neutralization in SARS-CoV-2 convalescent sera. Here we show that immunization with the SARS-CoV-2 RBD elicits a robust neutralizing antibody response in rodents, comparable to 100 µg/ml of ACE2-Ig, a potent SARS-CoV-2 entry inhibitor. Importantly, anti-sera from immunized animals did not mediate antibody-dependent enhancement (ADE) of S-protein-mediated entry under conditions in which Zika virus ADE was readily observed. These data suggest that an RBD-based vaccine for SARS-CoV-2 could be safe and effective.

## INTRODUCTION

Coronaviruses are enveloped single-stranded, positive-strand RNA viruses of the family *Corornaviridae* (Cui et al., 2019). They divide into four main subgroups: α, β, γ, and δ. At least seven coronaviruses infect humans: the α-coronaviruses HCoV-229E and HCoV-OC43, and the β-coronaviruses SARS-CoV (SARS-CoV-1), HCoV-NL63, CoV-HKU1, MERS-CoV, and the recently described SARS-CoV-2 (nCoV-19), a β-coronavirus closely related to human SARS-CoV-1 (79.0% nucleotide identity) and to SARS-CoV-like variants isolated in bats, including bat-SL-CoV-RaTG13 (96.2% nucleotide identity) (Lu et al., 2020; Menachery et al., 2015; Zhou et al., 2020). SARS-CoV-2 infection causes mild flu-like symptoms in many patients, but in many cases develops into an acute pulmonary syndrome (Chen et al., 2020; Zhou et al., 2020). SARS-CoV-1 causes severe acute respiratory syndrome (SARS), whereas disease associated with SARS-CoV-2 has been named COVID-19.

SARS-CoV-2, like SARS-CoV-1, requires expression of the cellular receptor ACE2 to infect cells (Hoffmann et al., 2020; Li et al., 2003; Walls et al., 2020). ACE2 is a carboxy metalloprotease that, among its other substrates, cleaves angiotensin I (angiotensin 1-10) and angiotensin II (angiotensin 1-8), respectively, to angiotensin (1-9) and angiotensin (1-7). In doing so it counteracts the vasoconstrictive activity of ACE1 (Hamming et al., 2007; Jia, 2016). Although the catalytic domains of ACE1 and ACE2 are similar, ACE1 inhibitors have no effect on the enzymatic activity ACE2. Moreover, the proteolytic activity of ACE2 does not contribute to SARS-CoV-1 or, presumably, SARS-CoV-2 entry processes (Li et al., 2003). However interference with ACE2 activity, through S-protein-promoted internalization and degradation, triggered release by cellular proteases, or loss of ACE2-expressing type II pneumocytes, may contribute to SARS and COVID-19 pathology. Although the physiologic role of ACE2 is complex, its activity is described as protective against acute respiratory distress syndrome (ARDS) caused by many agents (Imai et al., 2007). However administration of soluble recombinant ACE2, although well tolerated, did not attenuate acute lung injury in an early clinical study of ARDS unrelated to COVID-19(Khan et al., 2017).

Entry of SARS-CoV-2 into ACE2-expressing cells is mediated by its spike (S) protein (Hoffmann et al., 2020; Walls et al., 2020). The coronavirus S protein is a type I viral entry protein similar to influenza hemagglutinin and the HIV-1 envelope glycoprotein (Figure 1A and 1B) (Li, 2016). Like these latter entry proteins, the S protein is processed into two domains, S1 and S2 (Walls et al., 2020). S1 binds ACE2, whereas S2 anchors the S protein to the viral membrane. The SARS-CoV-2 S protein has an efficient furin cleavage site at its S1/S2 boundary, and this site is processed in virus-producing cells (Coutard et al., 2020). In contrast, the SARS-CoV-1 S1/S2 junction is cleaved by extracellular or target-cell proteases including TMPRSS2 and cathepsin L (Glowacka et al., 2011; Huang et al., 2006; Millet and Whittaker, 2015). Both S proteins require processing at a second site, S2’, within the S2 domain to mediate fusion of the viral and target cell membranes (Belouzard et al., 2009).

**Figure 1.**
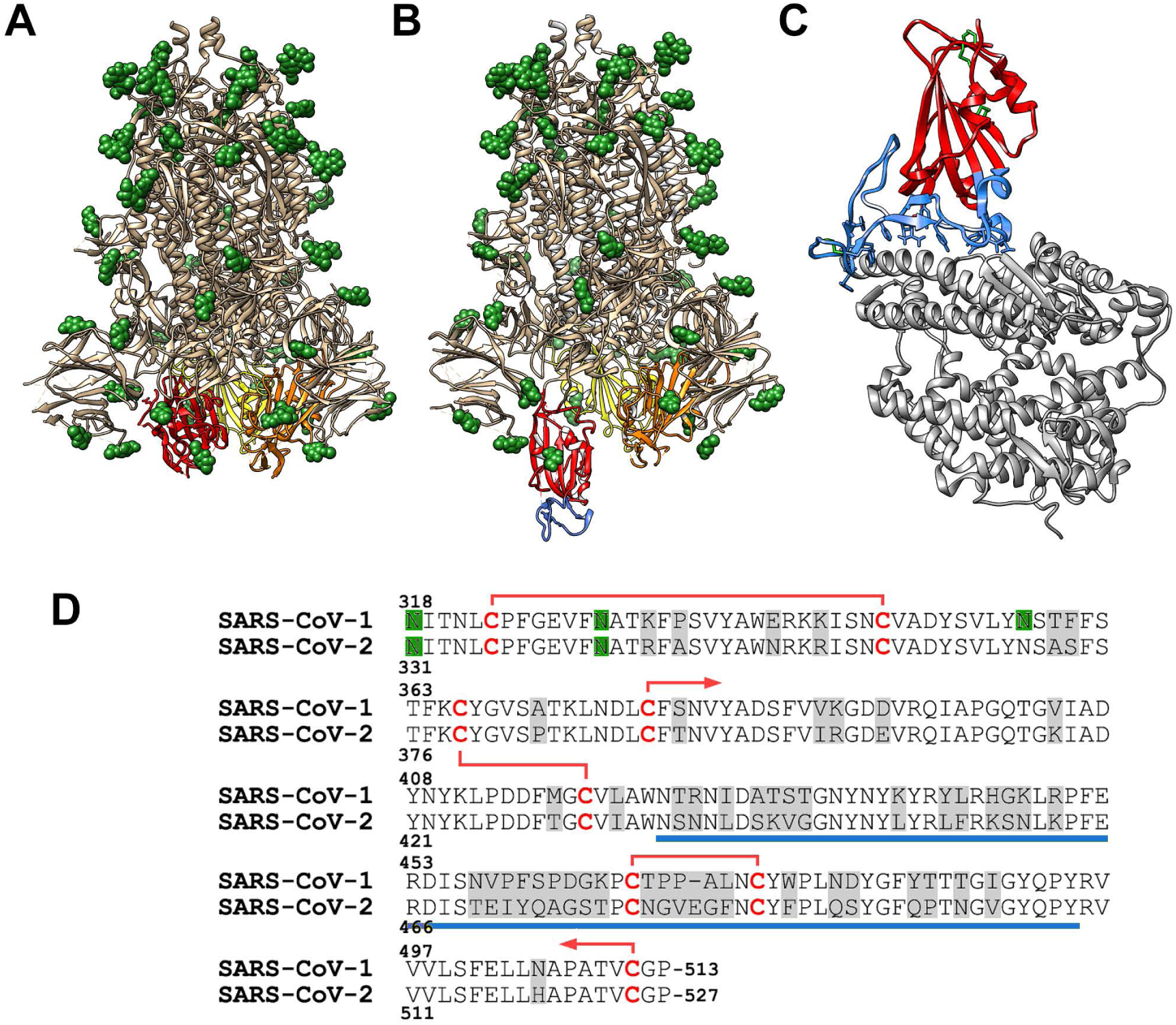
The SARS-CoV-2 RBD is a discrete domain of the S protein. The SARS-CoV-2 S protein trimer has been observed in at least two distinct pre-fusion conformations. (A) A symmetric state in which all three RBD domains (red, yellow, orange) are embedded symmetrically within the trimeric S-protein, and (B) a state in which at least one RBD domain (red) extends to interact with the cellular receptor ACE2. The S protein trimer is also modified by multiple N-linked glycosylations (green). The ACE2-binding motif of the RBD, termed the receptor-binding motif (RBM residues 437 to 508) is indicated in blue. (C) The RBD (red) is shown interacting with its receptor, human ACE2 (grey), and its four disulfide bonds are indicated in green. The RBM region is again indicated in blue. (D) The sequence of the SARS-CoV-1 and SARS-CoV-2 RBD motifs are shown aligned. Grey indicates differences between the two RBD, red and red lines indicate disulfide-linked cysteines, and the blue bar indicates the RBM region.

The receptor-binding domains (RBDs, also described as S^B^) of SARS-CoV-1 and SARS-CoV-2 directly bind ACE2 (Figure 1C) (Li et al., 2005; Walls et al., 2020; Wong et al., 2004; Wrapp et al., 2020). These RBDs are structurally and functionally distinct from the remainder of the S1 domain, and express and fold as independent domains (Wong et al., 2004). Both RBDs are highly stable and held together by four disulfide bonds. Structural studies of the SARS-CoV-2 RBD bound to ACE2 have identified a variable region, termed the receptor-binding motif (RBM), which directly engages ACE2 (Li et al., 2005). This region is divergent between SARS-CoV-1 and SARS-CoV-2 (Figure 1D) although both RBD bind ACE2 in the same orientation and rely on conserved, mostly aromatic, residues to engage this receptor. The divergence between the SARS-CoV-1 and SARS-CoV-2 RBM domains suggest that this region is subject to ongoing positive selection from the humoral response in various hosts. Despite this divergence, some antibodies, notably CR3022, bind both RBD domains (Tian et al., 2020).

Because the S protein is the dominant protein exposed on the virion, and because its activity can be impeded with antibodies, it is likely the major target of any SARS-CoV-2 vaccine. Soluble trimeric S proteins, including those stabilized through various mechanisms, have been tested as SARS-CoV-1 vaccines, and similar approaches are now being taken against SARS-CoV-2 (Chen et al., 2005; Walls et al., 2020; Wrapp et al., 2020). Another approach, immunizing with the RBD alone, has been shown to raise potent neutralizing antibodies against SARS-CoV-1 in rodents (He et al., 2006; He et al., 2004). Although the RBD presents fewer epitopes than the S-protein trimer, this approach may have key advantages. The RBD is easier to produce, and is less likely to elicit antibodies against poorly neutralizing but more immunogenic epitopes. In addition, antibody-dependent enhancement (ADE) contributes to the pathogenicity of feline coronaviruses (Huisman et al., 2009; Olsen et al., 1992; Weiss and Scott, 1981), and has been raised as a concern for vaccines against SARS-CoV-1 (Jaume et al., 2012; Liu et al., 2019; Luo et al., 2018; Wan et al., 2020; Wang et al., 2016; Wang et al., 2014; Yang et al., 2005; Yip et al., 2016)3. ADE contributes to the pathology of other viral diseases, notably those caused by flaviviruses, and in those cases, non-neutralizing antibodies promote more efficient ADE than those which also neutralize (Dejnirattisai et al., 2010; Halstead and O’Rourke, 1977; Takada and Kawaoka, 2003). Thus the RBD-elicited antibodies may mediate ADE less efficiently than those recognizing other S-protein epitopes because they are more likely to be neutralizing.

We accordingly investigated whether the SARS-CoV-2 RBD domain could raise efficient neutralizing antibodies and whether these antibodies could also mediate ADE. We show that this RBD raised potently neutralizing antisera in immunized rats and these sera did not mediate ADE under conditions where Zika-virus ADE was readily observed.

## RESULTS

The SARS-CoV-2 RBD, like that of SARS-CoV-1, is exposed in both known states of the S-protein trimer, namely a closed state where each RBD contacts symmetrically its analogues on the other protomer (Figure 1A), and an open state (Figure 1B) in which at least one RBD domain is extended to contact ACE2 (Figure 1C). We have previously shown that the SARS-CoV-1 RBD folds independently and expresses efficiently, and that an immunoadhesin form of this RBD neutralized S-protein mediated entry with a 50% inhibitor concentration (IC_50_) of ∼10 nM (Wong et al., 2004). This construct, RBD-Fc, also efficiently raised antibodies in mice capable of neutralizing SARS-CoV-1 variants with distinct RBD sequences (He et al., 2006; He et al., 2004). These previous data raised the possibility that an RBD-based SARS-CoV-2 vaccine could be effective against virus throughout the current COVID-19 pandemic.

To evaluate this possibility, we produced and purified the SARS-CoV-2 RBD (sequence shown in Figure 1D), fused to the Fc domain as an expedient for rapid purification. This RBD fusion protein was conjugated to a Keyhole limpet hemocyanin (KLH) carrier protein and mixed with the AS01 adjuvant formulation now used in at least two human vaccines. This antigen/adjuvant combination was injected intramuscularly into four female Sprague-Dawley rats with a schedule of seven increasing (2.5-fold) doses, one each day, ultimately administering a total of 500 µg of the SARS-CoV RBD-Fc. Thirty days after the first administration, RBD fused to a 4-amino acid C-tag was purified with a C-tag affinity column, again conjugated to KLH, and the immunization regimen was repeated. Again a total of 500 µg of RBD was administered.

Blood was harvested from each of the four rats (R15, R16, R17, R18) immediately before inoculation (day 0), and at 5-day intervals starting at day 10 after the first inoculation. Serial dilutions of day-0 and day-40 sera were measured for their ability to neutralize retroviruses pseudotyped with the SARS-CoV-2 S protein (SARS2-PV). To provide a consistent metric for neutralization potency, these sera were also compared with a mixture of all four day-0 preimmune sera, further combined with an immunoadhesin form of ACE2 (ACE2-Ig) at concentrations in sera of 10, 100, and 1000 µg/ml before dilution. As anticipated, some baseline inhibition could be observed in heat-inactivated rat sera (grey lines in Figure 2A). However day-40 serum from each rat, obtained after two sets of immunizations, potently neutralized SARS2-PV entry with an efficiency comparable to or greater than day-0 presera mixed with ACE2-Ig at a 100 µg/ml concentration (Figure 2A and 2B). Indeed day-40 serum from one of four rats, R16, neutralized as efficiently as 1 mg/ml of ACE2-Ig mixed into preimmune serum. For context, the concentration of IgG in rat serum in approximately 12-14 mg/ml (Medesan et al., 1998). ACE2-Ig itself, in the context of rat presera, appears to have an IC_50_ of well under 1 µg/ml (10 nM), as a 1620-fold dilution of 1000 µg/ml ACE2 in preimmune sera neutralized more than 50% of SARS2-PV entry (Figure 2B). This ACE2-Ig inhibition is greater than what we observed with SARS-CoV-1 (Moore et al., 2004), consistent with ACE2’s higher affinity for the SARS-CoV-2 RBD (Walls et al., 2020; Wrapp et al., 2020). We conclude that the SARS-CoV-2 RBD can elicit a neutralizing response in vaccinated rats comparable to a 100 µg/ml (1 µM) concentration of an inhibitor with a 1 nM IC_50_.

**Figure 2.**
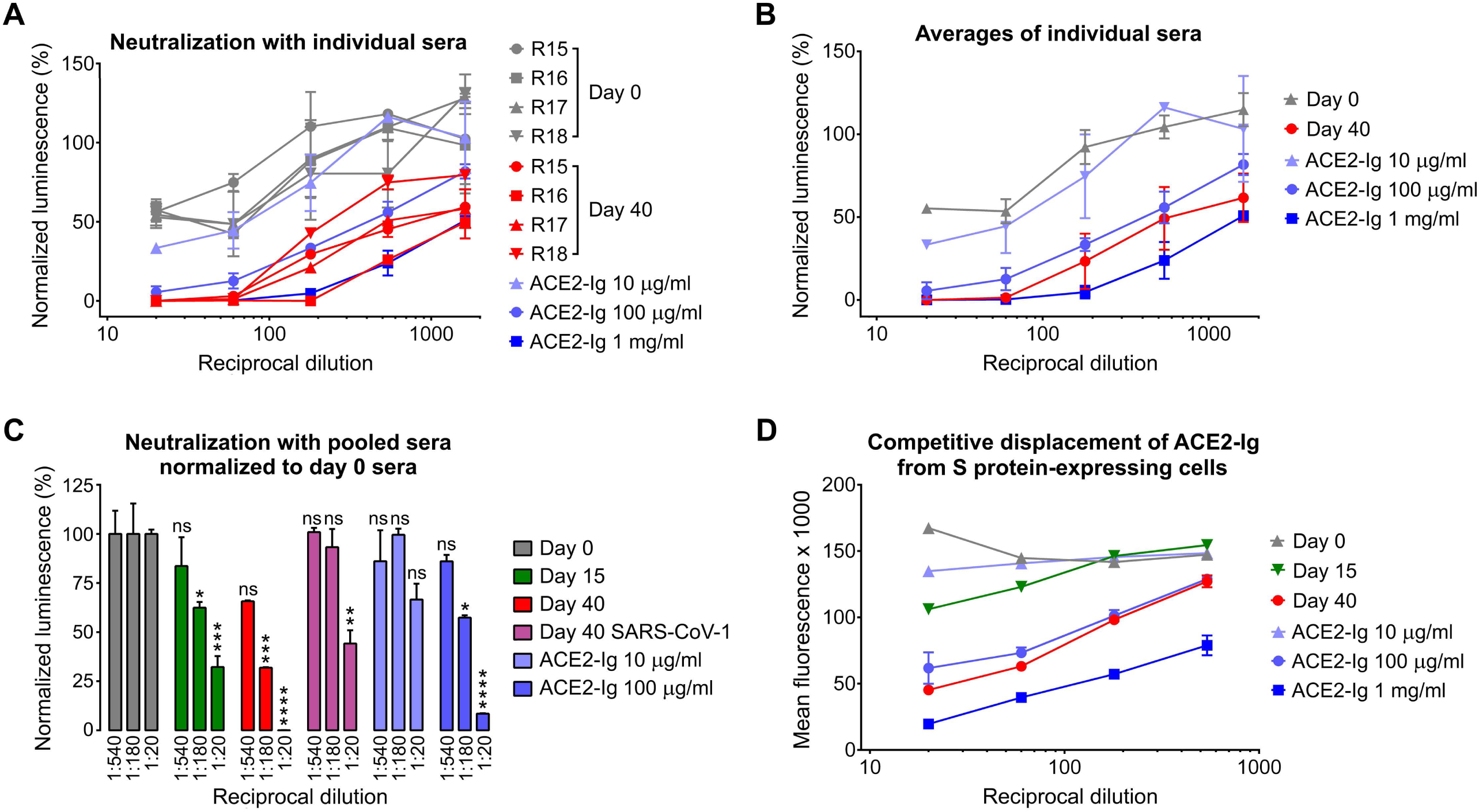
Immunization with the SARS-CoV-2 RBD elicits potently neutralizing antibodies. Four female Sprague Dawley rats (R15, R16, R17, R18) were immunized with two sets of escalating doses of RBD conjugated to keyhole limpet hemocyanin. (A) The indicated dilutions of preimmune sera (day 0, gray) were compared to dilutions of sera harvested from immunized rats at day 40, and to the same dilutions of preimmune sera mixed to achieve the indicated ACE2-Ig concentrations before dilution. Each serum and serum-ACE2-Ig mixture was compared for its ability to neutralize S-protein-pseudotyped retroviruses (SARS2-PV), by measuring the activity of a firefly-luciferease reporter expressed by these pseudoviruses. Figure shows entry of SARS2-PV as a percentage of that observed without added rat serum. Error bars indicate the range of two neutralization studies. (B) The data from each rat of panel A are averaged for clarity. Error bars indicate s.e.m., with each rat considered a different experiment. Differences between day-0 and day-40 serum are significant at all dilutions (*P* < 0.001; two-way ANOVA). (C) Sera from each rat was pooled before characterization, and SARS2-PV entry was measured at the indicated dilutions. Day-0 preimmune sera, day-15 sera, and day-40 sera were compared to day-0 sera mixed with ACE2-Ig to achieve the indicated concentrations before dilution, and characterized for their ability to neutralize SARS2-PV. Day 40 sera was also tested for its ability to neutralize SARS-CoV-1 pseudoviruses (SARS1-PV) as indicated. Values shown indicate percentage of pseudovirus entry observed with the same dilution of day-0 preimmune sears. Significant differences compared to day-0 serum at each dilution are indicated above the bar (* indicates *P* < 0.05; ** indicates *P* < 0.01; *** indicates *P* < 0.005; **** indicates *P* < 0.001; ns indicates *P* > 0.05; two-way ANOVA). Error bars indicate range of two entry assays. (D) Pooled sera and pooled preimmune sera mixed with the indicated concentrations of ACE2-Ig were further combined with an ACE2-Ig variant bearing a rabbit-derived Fc domain (ACE2-rIg). Binding of the ACE2-rIg was monitored with an anti-rabbit Fc secondary antibody as determined by flow cytometry. Error bars indicate the range of two such measurements. Differences between day-0 and day-40 serum are significant (*P* < 0.001; two-way ANOVA) at all dilutions.

To further characterize the effects of the two successive sets of immunizations, we compared day-0, day-15, and day-40 sera by pooling these sera from all rats, and also compared it with pooled day-0 serum mixed with 10 and 100 µg/ml ACE2-Ig, diluted as indicated (Figure 1C). Again day-40 serum potently neutralized S-protein-mediated entry, with significantly greater efficiency than day-0 serum mixed with 100 µg/ml ACE2-Ig. At lower dilutions (1:20 and 1:60) day-15 serum neutralized more efficiently than day-0 serum, somewhere between 10 and 100 µg/ml ACE2 in preserum. However day-40 serum did not efficiently neutralize retroviruses pseudotyped with SARS-CoV-1 S protein (SARS1-PV), except at the lowest (1:20) dilution, excluding the possibility that immunization with a single RBD could protect against SARS-like viruses with divergent RBD. Finally, to confirm that sera from vaccinated rats neutralized SARS2-PV by recognizing the SARS-CoV-2 S protein, we used each pooled serum to displace an ACE2-Ig variant with a rabbit Fc (ACE2-rIg) domain from cells expressing the SARS-CoV-2 S protein (Figure 2D). The ability of each pooled serum to displace ACE2-rIg correlated with its ability to neutralize SARS2-PV. Specifically, day-40 sera displaced ACE2-rIg more efficiently than preimmune sera containing 100 µg/ml ACE2-Ig, and day-15 sera displaced ACE2-rIg more efficiently than preimmune sera containing 10 µg/ml ACE2-Ig. Thus immunization with SARS-CoV-2 RBD elicits antibodies that potently neutralize S-protein mediated entry by directly binding the S protein.

One concern associated with coronavirus vaccines is the possibility that anti-S-protein antibodies could increase the efficiency of infection of cells, such as alveolar macrophages, expressing Fc receptors, for example FcγRI (CD64) or FcγRII (CD32). This undesirable antibody-dependent enhancement (ADE) has been well characterized in tissue-culture studies of several flaviviruses including Zika virus (ZIKV) and dengue virus (Shim et al., 2019). To evaluate this possibility for SARS-CoV-2, SARS2-PV were mixed with pooled day-0 or day-40 serum at the indicated serial dilutions, and the resulting virus/sera mixtures were incubated with HEK293T cells transfected to express rat FcγRI. These cells did not express ACE2 and no infection was observed with day-0 preimmune sera nor day-40 immune sera (Figure 3A). To verify that these cells were capable of mediating ADE, rat anti-ZIKV antisera generated by two different rats and pooled, or day-0 preimmune sera was incubated at the same dilutions with ZIKV virus-like particles (VLP). In contrast to the absence of effect of day-40 anti-RBD anti-sera on SARS2-PV, anti-ZIKV antisera at the same concentrations promoted robust ADE (Figure 3B). ADE activity peaked at approximately a 1/2000 dilution, consistent with competition between ADE and neutralizing activities of these antisera.

**Figure 3.**
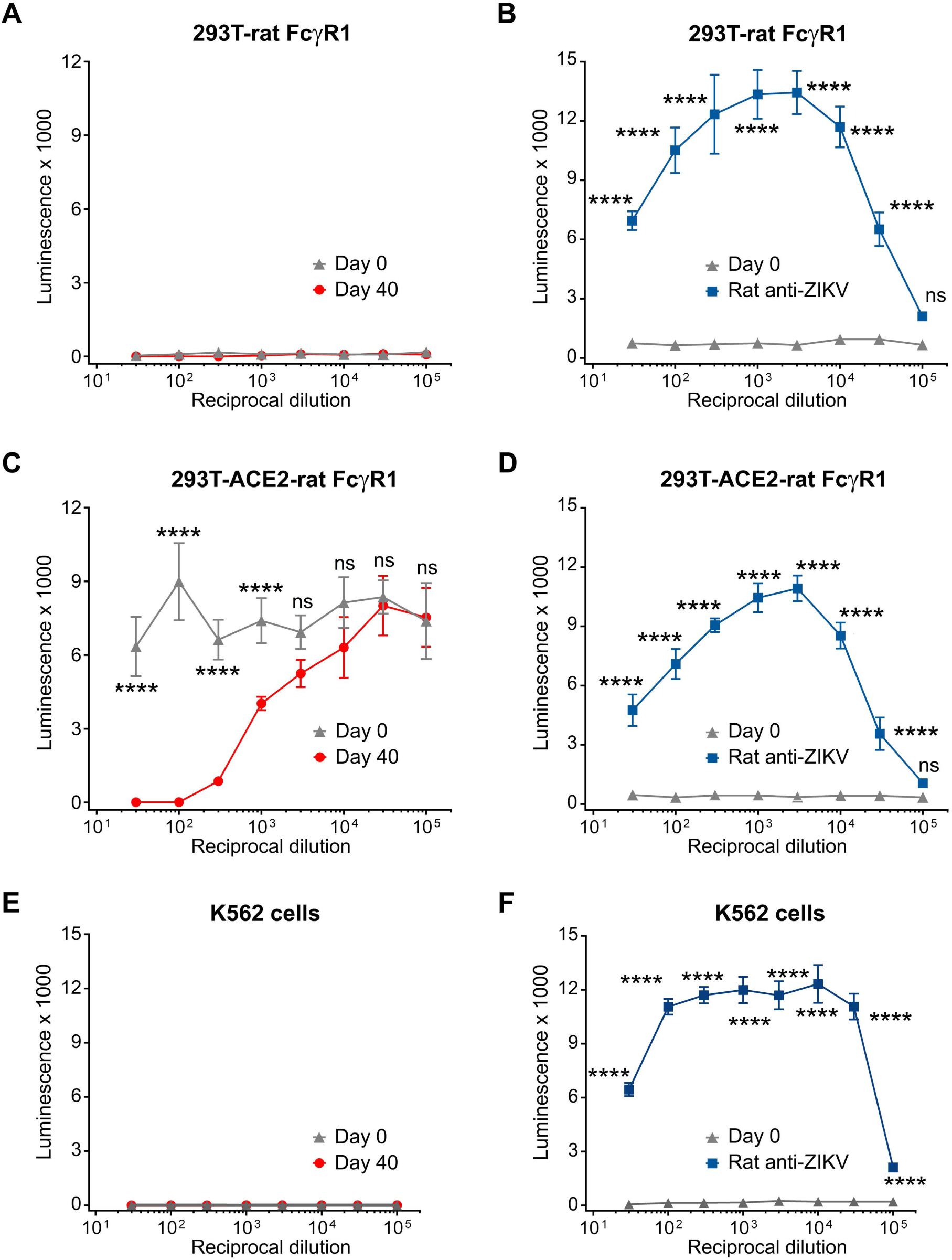
Anti-RBD antisera does not mediate antibody-dependent enhancement of SARS-CoV-2 S protein-mediated entry. (A) HEK293T cells were transfected to express rat FcγRI and infected with SARS2-PV in the presence of the indicated dilutions of preimmune day 0 or immune day 40 antisera. (B) Experiments similar to those in panel A except that ZIKV VLP and anti-ZIKV antisera, also elicited in Sprague Dawley rats, were compared to preimmune sera. (C, D) Experiments identical to those in panel A and B respectively except that HEK239T cells stably expressing human ACE2 were again transfected to express rat FcγRI before incubation with SARS2-PV (C) or ZIKV-VLP (D). (E, F) Experiments identical to A and B, respectively, except that the pro-monocytic K562 cell line was incubated with SARS2-PV (E) or ZIKV-VLP (F). Error bars in panels A-F indicate the range of two experiments run in parallel. Significant differences with day-0 preimmune sera are indicated (**** indicates *P*<0.001 by two-way ANOVA; ns indicates not significant).

To determine if ADE could be observed in the presence of the SARS-CoV-2 receptor ACE2, HEK293T cells stably expressing ACE2 were transfected to express rat FcγRI. Day-0 preimmune sera had no detectable impact on SARS2-PV infection, whereas day-40 serum efficiently neutralized this infection (Figure 3C). However, Day-40 serum did not promote infection at serum dilutions at which anti-ZIKV antiserum again promoted robust ADE of ZIKV VLP (Figure 3D). Thus anti-RBD anti-serum did not promote ADE at serum dilutions and under conditions in which ZIKV ADE could be easily observed. Finally we confirmed the results shown in Figure 3A and B using K562 cells that endogenously express FcγRII (Chiofalo et al., 1988). Again, day-40 sera did not enhance entry SARS-PV entry, but rat anti-ZIKV antisera again robustly promoted ZIKV-VLP infection (Figure 3E and 3F). Collectively the data of Figure 3 suggest that antibodies elicited by an RBD-based vaccine are likely to neutralize more efficiently than they would mediate ADE.

## DISCUSSION

A vaccine against SARS-CoV-2 should, for several reasons, be easier to generate than those against many other viruses (Gralinski and Baric, 2015; Li, 2016). First, coronaviruses have exceptionally large genomes compared to other RNA viruses, and, to avoid error catastrophe, their viral polymerase has acquired a proof-reading function. Thus any individual gene, for example that of the S protein, is likely to retain its original sequence over multiple replication cycles. Second, coronaviruses in general, and clearly SARS-CoV-2 in particular, transmits to a new host more rapidly than an adaptive immune response can emerge. A likely consequence of this strategy is that one of its most critical epitopes, namely its RBD, is exposed on the virion (Li, 2016; Walls et al., 2020; Wong et al., 2004; Wrapp et al., 2020), favoring transmission efficiency over antibody resistance. Finally, the stability and compactness of the SARS-CoV-1 and SARS-CoV-2 RBD means that it can be easily manufactured and presented to the immune system using many production technologies, presentation scaffolds, and delivery systems (Wong et al., 2004).

Many efforts are now underway to develop vaccines based on stabilized soluble trimeric forms of the S protein ectodomain (Wrapp et al., 2020). It is at this moment unclear whether these S-protein trimers, RBDs, or some parallel or serial combination thereof, will make better antigens. Recent studies suggest that neutralization activity of patient sera correlates with its RBD recognition, and that most neutralizing antibodies bind the RBD (To et al., 2020). It is possible that the remainder of the S protein presents non-neutralizing epitopes that are more immunogenic than the RBD, and thus the RBD by itself may elicit antisera that are more neutralizing. However, other antibody effector functions, including antibody-dependent cell-mediated cytotoxicity (ADCC) and complement fixation, are more efficiently activated when multiple antibodies bind their target simultaneously, and these non-neutralizing activities can play important roles in viral control. If these activities contribute significantly to SARS-CoV-2 prevention, the S-protein trimer may make a better antigen than the RBD. Alternatively, an optimized combination of both antigens may focus the immune response to a critical neutralizing epitope while also promoting more efficient antibody effector functions. A trimer-prime, RBD-boost strategy may even be necessary. For example, the first available vaccines expressing soluble trimers may be insufficiently effective in older and immunocompromised individuals. A subsequent RBD-based boost might improve protection in these individuals.

However, development of a safe coronavirus vaccine is complicated by the possibility that such a vaccine may contribute to COVID-19 pathology. There are now multiple examples in the literature whereby a SARS-CoV-1 vaccine can have a deleterious effect in non-human recipients (Jaume et al., 2012) (Luo et al., 2018; Wang et al., 2016; Yip et al., 2016), and SARS-CoV-1 ADE has been observed by several groups in cell culture studies (Jaume et al., 2011; Liu et al., 2019; Wan et al., 2020; Wang et al., 2014; Yang et al., 2005; Yip et al., 2014). ADE is more obvious in feline coronavirus, where it results in a shift in tropism and increase in disease severity (Huisman et al., 1998; Huisman et al., 2009; Weiss and Scott, 1981). Although ADE could theoretically be mediated by multiple mechanisms, the standard paradigm is provided by flaviviruses, most notably by dengue virus (Takada and Kawaoka, 2003). It is well appreciated that a second dengue virus infection with a different serotype can result in a hemorrhagic fever. In addition, a dengue-virus vaccine was reported to increase hospitalizations of vaccinees after their first dengue-virus exposure (Flipse and Smit, 2015; Tsai et al., 2017). In both cases, the underlying mechanism is presumed to be enhanced Fc-receptor-mediated internalization of virions bound to antibodies.

As observed with flaviviruses, there is a qualitative difference between ADE mediated by neutralizing and by non-neutralizing antibodies (Dejnirattisai et al., 2010; Shim et al., 2019; Takada et al., 2003). In the case of neutralizing antibodies, ADE is a consequence of insufficient coating of the virion surface, due to lower affinity or concentration of the antibody. If concentrations or affinity is raised, neutralization dominates. In contrast, ADE mediated by non-neutralizing antibodies increases with increasing concentration. We therefore speculated that it would be difficult of observe ADE with rat antisera targeting the SARS-CoV-2 RBD, a key neutralizing epitope. Indeed, although rat anti-Zika virus antisera mediated robust ADE of ZIKV in two different Fc receptor-expressing cell lines, no ADE of S-protein pseudoviruses was observed with anti-RBD anti-sera, whether or not the SARS-CoV-2 receptor ACE2 was present. These data suggest that, at least for RBD-based vaccines, ADE is likely to be less of a concern than with vaccines against flaviviruses such as ZIKV or dengue viruses. However they do not exclude ADE mediated by other S-protein epitopes or other mechanisms by which a vaccine can be deleterious.

A key remaining question is whether these studies, conducted in rodents, imply that an RBD-based vaccine could be effective in humans. Here experimental details become important. We immunized rats with an adjuvant system (AS01) already in human use, notably for vaccines for shingles (Shingrix™) and malaria (Mosquirix™). The RBD used here was conjugated to a carrier protein, Keyhole limpet hemocyanin, proven safe over decades in humans (Swaminathan et al., 2014). However KLH can be problematic for those with shellfish allergies, and alternate scaffolds for presenting the RBD may be necessary. Rats were immunized with two sets of escalating inoculations, each administered over seven days. This strategy has been shown to maximize immune responses against HIV-1 antigens (Tam et al., 2016), but it may not be a practical means of addressing the current pandemic with a subunit vaccine. That said, the escalating dosage strategy used here may more accurately reflect the kinetics of an mRNA-delivered vaccine, and indeed we would argue that an RBD antigen would be a preferable alternative, or at least a useful complement, to mRNA expression of the S-protein trimer.

However, it remains to be determined whether a more limited number of injections of an RBD subunit vaccine would raise a protective response. A key point relevant to this question is that the neutralizing activity elicited in these studies is likely much greater than would be necessary to prevent a new SARS-CoV-2 infection. Rat antisera on average neutralized more efficiently than naive sera bearing 100 µg/ml of ACE2-Ig, which itself neutralizes with an IC_50_ of less than 1 µg/ml, consistent with its activity against infectious SARS-CoV-1 (Moore et al., 2004). To provide some context for these numbers, eCD4-Ig, an immunoadhesin that emulates HIV-1 viral receptors and neutralizes with neutralization potency similar to that of ACE2-Ig, affords robust protection against high-dose intravenous challenges with an HIV-1-like virus when it is present in serum in the range of 5-10 µg/ml (Gardner et al., 2019; Gardner et al., 2015). Thus it is possible that a less robust response could still be protective. Perhaps as important as these quantitative considerations, our data show that the RBD alone is sufficient to elicit a potent neutralizing antibody response. It remains to be determined whether trimer-based vaccines, or trimer/RBD combinations, can be more effective, and whether they carry additional safety concerns.

In short, our data suggest that an RBD vaccine could protect many individuals from SARS-CoV-2 and that it is unlikely to promote infection through conventional ADE mechanisms. Given the exposure of this key neutralizing epitope, the ease with which this compact domain can be produced and presented, and its clear immunogenicity, we propose that the RBD be considered as a key component of any SARS-CoV-2 vaccine strategy.

## AUTHOR CONTRIBUTIONS

B.D.Q., H.C., and M.F. conceived the study, B.D.Q., H.M., L.Z, and H.C. designed experiments, B.D.Q, H.M., L.Z., Y.G, W.H, and A.O. performed experiments. M.S.P., G.L., W. L., and G.Z. provided critical reagents and advice. M.F. wrote the manuscript with critical assistance from H.C.

## MATERIALS AND METHODS

### Contact for Reagent and Resource Sharing

Further information and requests for resources and reagents should be directed to and will be fulfilled by Michael Farzan.

### Production of SARS-CoV-1 and SARS-CoV-2 pseudoviruses and Zika-virus virus-like particles

Retroviruses pseudotyped with the SARS-CoV-2 or SARS-CoV-1 S proteins (SARS2-PV, SARS1-PV) were produced as previously described (Moore et al., 2004) with modest modifications as described. HEK293T cells were transfected by polyethylenimine (PEI) transfection at a ratio of 5:5:1 with a plasmid encoding murine leukemia virus (MLV) gag/pol proteins, a retroviral vector pQCXIX expressing firefly luciferase, and a plasmid expressing the spike protein of SARS-CoV-1 (GenBank AY271119) or SARS-CoV-2 (GenBank YP_009724390). Cells were washed 6 hours later, and the culture supernatant containing pseudoviruses was harvested at 48-72 hours post transfection. Zika virus virus-like particles (ZIKV-VLP) were produced by transfecting HEK293T cells by the calcium phosphate transfection method with a ZIKV replicon (strain FS13025, GenBank KU955593.1), whose expression is controlled by tetracyclin, a plasmid encoding ZIKV capsid, prM, and E proteins (strain FSS13025, GenBank KU955593.1), and the pTet-On plasmid expressing a reverse Tet-responsive transcriptional activator (rtTA) at a ratio of 2:1:1. Cells were washed 6 h later and replenished with fresh media containing 1 μg/ml doxycycline. The VLP-containing culture supernatant was harvested 48 h post transfection. ZIKV replicon was generated by replacing the region spanning 39^th^ through 763^rd^ amino acids of the polyprotein of a ZIKV molecular clone we previously generated (Zhang et al., 2018) with Renilla luciferase with the 2A self-cleaving peptide fused at its C-terminus. This construct contains the tetracyclin-responsive P_tight_ promoter that drives ZIKV RNA transcription. The pseudovirus- or VLP-containing culture supernatants were cleared by 0.45 µm filtration. SARS2-PV and ZIKV-VLP titers were assessed by RT-qPCR targeting the CMV promoter in the retroviral vector pQCXIX and ZIKV NS3 gene, respectively. In some cases, clarified pseudovirus and VLP stocks were stored at -80°C for long-term storage and reuse.

### Generation of human ACE2 expressing cell lines

HEK293T cells expressing human ACE2 (hACE2) were generated by transduction with murine leukemia virus (MLV) pseudotyped with the vesicular stomatitis virus G protein and expressing myc-hACE2-c9, as previously described (Wicht et al., 2014). Briefly, HEK293T cells were co-transfected by PEI with three plasmids, pMLV-gag-pol, pCAGGS-VSV-G and pQCXIP-myc-hACE2-c9 at a ratio of 3:2:1, and medium was refreshed after overnight incubation of transfection mix. The supernatant with produced virus was harvested 72-hours post transfection and clarified by passing through 0.45μm filter. 293T-hACE2 cells were selected and maintained with medium containing puromycin (Sigma). hACE2 expression was confirmed by SARS1-PV and SARS2-PV entry assays and by immunofluorescence staining using mouse monoclonal antibody recognizing c-Myc.

### Protein Production

Expi293 cells (Thermo-Fisher) were transiently transfected using FectoPRO (Polyplus) with plasmids encoding SARS-CoV2 RBD with a human or rabbit-Fc fusion or a C-terminal C-tag (-EPEA). After 5 days in shaker culture, media were collected and cleared of debris for 10 min at 1,500 g nd filtered using 0.45-µm flasks (Nalgene). Proteins were isolated using MabSelect Sure (GE Lifesciences) or CaptureSelect C-tagXL (Thermo-Fisher) columns according to the manufacturers’ instructions. Eluates were buffer exchanged with PBS and concentrated using Amicon ultra filtration devices (Millipore Sigma) and stored at 4°C before use..

### Immunizations and sera collection

All rats used in these studies were handled and maintained in accordance with NIH guidelines and approved by Institutional Animal Care and Use Committee (IACUC) of Scripps Research (Protocol 18-025). Female Sprague Dawley rats were immunized with incremental increasing doses of antigen over seven days starting at day 0, and boosted with a similar regimen at day 30. Rats were inoculated in the first set with the SARS-CoV-2 RBD fused to the Fc domain of human IgG1, and in the second set with the RBD fused to a four amino-acid C-tag. In both cases RBD fusions were conjugated at a 1:1 ratio to Mariculture keyhole limpet hemocyanin (mcKLH, Thermo-Fischer Peirce) by 1-ethyl-3-(3-dimethylaminopropyl)carbodiimide hydrochloride (EDC, Thermo-Fischer Peirce) according to manufactures protocols. Each set of seven injections were performed in the following manner. RBD-KLH conjugates were administered intramuscularly into the rear quadriceps. Inoculations were initiated with 2.2 µg RBD-KLH (equivalent to 1.1 µg RBD antigen) adjuvanted with 0.1 µg MPLA and 0.1 µg QuilA, and this inoculum was increased by 2.55 fold for each of the next six days, administering a total of 500 µg RBD-Ig or RBD-C-tag fusion protein, and 40 µg of each adjuvant component. Sera were collected before inoculation (day 0 preimmune sera) and every five days starting on 10^th^ day after the first injection. All sera was heat-inactivated for 30 minutes at 56°C and stored at -80°C for reuse.

### Neutralization studies of SARS-CoV-1 and SARS-CoV-2 pseudoviruses

Individual sera or pooled sera collected at day 0 (pre-immune sera), day 15, and day 40 after the first inoculation were serially diluted in DMEM. In some cases, day 0 sera were mixed with ACE2-Ig to a concentration of 10, 100, or 1000 µg/ml before dilution, and then diluted in the same manner. Individual, pooled, or pooled-ACE2-Ig sera was mixed with SARS1- or SARS2-PV and incubated at 37°C for one hour. One hour later, 10^4^ ACE2-239T cells were added along with DEAE-Dextran (final concentration 5 µg/ml), and media was exchanged 6 hours later with fresh media without rat sera. At least two independently mixed replicates were measured for each experiment. Firefly luciferase activity was measured (Britelight) 48 hours post-infection. All neutralization studies were performed at least twice with similar results.

### Competitive displacement of ACE2-Ig from S-protein expressing cells

Serial dilutions of pooled sera or pooled pre-immune sera mixed with ACE2-Ig (with the human Fc domain) at initial concentrations of 10, 100 and 1000 µg/mL were then mixed with 1 µg/ml ACE2-rIg (with a rabbit Fc domain). Independent mixtures were made for each replicate. Pooled sera and sera mixtures were used to stain HEK293T cells transfected by the jetPRIME Transfection Reagent (Polyplus) to express the full-length SARS-CoV2 S protein. Specifically, 10^5^ per cells per well were placed in a 96-well V-bottom plate and incubated for 45 minutes at 4°C with 100 μl of the serially diluted sera doped with ACE2-rIg. After washing, cells were stained with anti-rabbit IgG-Alexa647 antibody for 45 minutes at 4°C, and mean fluorescence intensities were measured for each well by flow cytometry.

### Measurement of antibody-dependent enhancement

The ability of anti-SARS-CoV-2 RBD immune plasma to mediate antibody-dependent enhancement (ADE) was measured using HEK293T cells or HEK293T cells stably expressing human ACE2 (293T-ACE2 cells), transfected using the calcium phosphate transfection method to express the rat ortholog of FcγRI (CD64). The human monocytic cell line K562 (ATCC CCL-243), which endogenously expresses FcγRII, was also used for ADE assays. The RBD immune sera, collected from four different rats at day 40 after the first immunization, were mixed at an equal ratio, as was preimmune sera obtained at day 0 from the same rats. As a positive control, sera from ZIKV-infected rats (rat #13 and #15, distinct from the similarly numbered RBD-inoculated rats), was also mixed at an equal ratio. Immune and pre-immune serum samples were heat inactivated for 30 min at 56°C and serially diluted in DMEM containing 10% heat-inactivated FBS. SARS2-PV or ZIKV-VLP in 50 µl was preincubated for 1 h at 37°C with 50 µl of diluted sera and added to the indicated cells plated on the 96 well plates. Two days later, infection levels were assessed using Luc-Pair Firefly Luciferase HS Assay Kit (Genocopia) for SARS-PV and Luc-Pair Renilla Luciferase HS Assay Kit (Genocopia) for ZIKV-VLP.

### Statistical analyses

The statistical significance of differences between preimmune and immune sera, or ACE2-Ig, in their abilities to neutralize, bind the S protein, and mediate ADE was analyzed by two-way ANOVA calculated using GraphPad Prism 8.0. Differences were considered significant at *P* < 0.05.

## Notes

### Competing Interest Statement

The authors have declared no competing interest.

